# A universal framework for detecting cis-regulatory diversity in DNA regulatory regions

**DOI:** 10.1101/2020.10.26.354522

**Authors:** Anushua Biswas, Leelavati Narlikar

## Abstract

High-throughput sequencing-based assays measure different biochemical activities pertaining to gene regulation, genome-wide. These activities include protein-DNA binding, enhancer-activity, open chromatin, and more. A major goal is to understand underlying sequence components, or motifs, that can explain the measured activity. It is usually not one motif, but a combination of motifs bound by cooperatively acting proteins that confers activity to such regions. Furthermore, although having a single type of activity, the regions can still be diverse, governed by different combinations of proteins/motifs. Current approaches do not take into account this issue of combinatorial diversity. We present a new statistical framework cisDiversity, which models regions as diverse modules characterized by combinations of motifs, while simultaneously learning the motifs themselves. We show that ChIP-seq data for the CTCF protein in fly contains diverse sequence structures, with most direct CTCF-binding sites situated far from promoters, giving insights into its co-factors and potential role in looping. Human CTCF-bound regions, on the other hand, have a different architecture. Because cisDiversity does not rely on knowledge of motifs, modules, cell-type, or organism, it is general enough to be applied to regions reported by most high-throughput assays. Indeed, enhancer predictions resulting from different assays—GRO-cap, STARR-seq, and those measuring chromatin structure—show distinct modules and combinations of TF binding sites, some specific to the assay. No module occurs universally in all enhancer-assays. Finally, analysis of accessible chromatin suggests that regions open in one cell-state encode information about future states, with certain modules staying open and others closing down later. The code is freely available at https://github.com/NarlikarLab/cisDIVERSITY.

## 1 Introduction

High-throughput sequencing technologies are routinely used to map multiple types of biochemical activities occurring across the genome. Examples include protein-DNA binding events ^1,2^, open chromatin regions ^3–5^, interacting chromatin domains ^6,7^, active transcription start sites (TSSs) ^8^, and many more ^9^. A major goal of these efforts is to understand what part of the underlying sequence might be driving that particular activity. Now, although the measured activity might be of a specific nature, the same sequence signature may not be responsible for it at all locations. Consider, for example, an experiment such as ATAC-seq or DNase-seq, which identifies open chromatin regions. The reason behind the accessibility of a region may be one of several: it may be an active promoter, or an enhancer, or an insulator or even a matrix-attachment region. Naturally, then, the pertinent sequence components in those regions will also be different. In some cases the heterogeneity is less obvious, but present, all the same. For instance, while the primary objective of a high-throughput chromatin immunoprecipitation (ChIP) experiment is to identify regions bound by a specific transcription factor (TF), in reality the experiment reports a miscellaneous set of genomic regions: those making direct contact with the TF, those indirectly bound to the TF via an intermediate, those where the TF binds along with a co-factor, and perhaps, those that are simply proximal to the TF in 3D space ^10^.

While the existence of such an assortment of regions is well-accepted in most high-throughput experiments, methods used to learn the regulatory architecture at these regions do not effectively account for it. Reported regions are generally analyzed by identifying individual overrepresented motifs, by sequentially searching for motifs one after the other, either from a known database or de novo. This can fail in certain situations. Motifs may be missed because they are present only in a small set of regions and therefore not statistically overrepresented in the entire set. A few recent methods do account for this by posing this as a clustering problem, with each cluster of sequences being dominated by a potentially different motif ^11,12^. However, these approaches do not take into account combinations of motifs, which may be critical to drive the biochemical activity at the region. Additionally, there may be multiple distinct motif-combinations across the reported regions, with each combination explaining a fraction of the regions.

Here we propose a new method called cisDiversity, which attempts to explain the whole set of the reported regions in terms of motifs and their combinations, all computed de novo. We take inspiration from topic modeling in computer science, where the goal is to cluster documents (here: DNA regions) into different topics (here: functions/modules), based on word frequencies (here: DNA motifs). DNA regions come from a categorical distribution over modules, and each module is a product of Bernoulli distributions over presence/absence of motifs. In contrast to typical topic modeling where the words are established and parsed a priori, here the motifs are also unknown and learned along with the modules. The only input is the set of sequences obtained from the experiment.

We apply cisDiversity to a range of different high-throughput datasets: ChIP-seq, eRNAs, STARR-seq, ATAC-seq, and DNase-seq. In each case cisDiversity identifies modules composed of different combinations of de novo motifs. These modules appear to have different characteristics, although they belong to the same experimental dataset, showing the utility of cisDiversity.

## 2 Results

### 2.1 A sequence model for representing cis-regulatory diversity

A high-throughput sequencing experiment typically identifies a set ***X*** of genomic regions that are enriched with a specific biochemical activity. We assume that *r* different regulatory modules may be responsible for the activity and each module is defined by the presence or absence of *m* motifs. More specifically, each module is modeled as a product of *m* Bernoulli distributions corresponding to the presence or absence of each motif. Different modules will have different Bernoulli distributions over the same motifs. A motif is modeled with the standard position weight matrix (PWM) ^13^. Figure 1A shows an instance of a simulated dataset of a 1000 sequences, where five motifs were planted from the JASPAR database ^14^, with specific Bernoulli distributions across three modules. For instance, motif 1 is present in all sequences of module 2, in a fifth of sequences of module 1, and never in any of module 3. On the other hand, motif 5 is present in all sequences of module 3, but because the module consists of only about 40 sequences, overall motif 5 occurs less frequently in the data. The aim of cisDiversity is to learn the *m* motif parameters, the *r* × *m* Bernoulli distributions, and the sequences that belong to each of the *r* modules.

**Figure 1:**
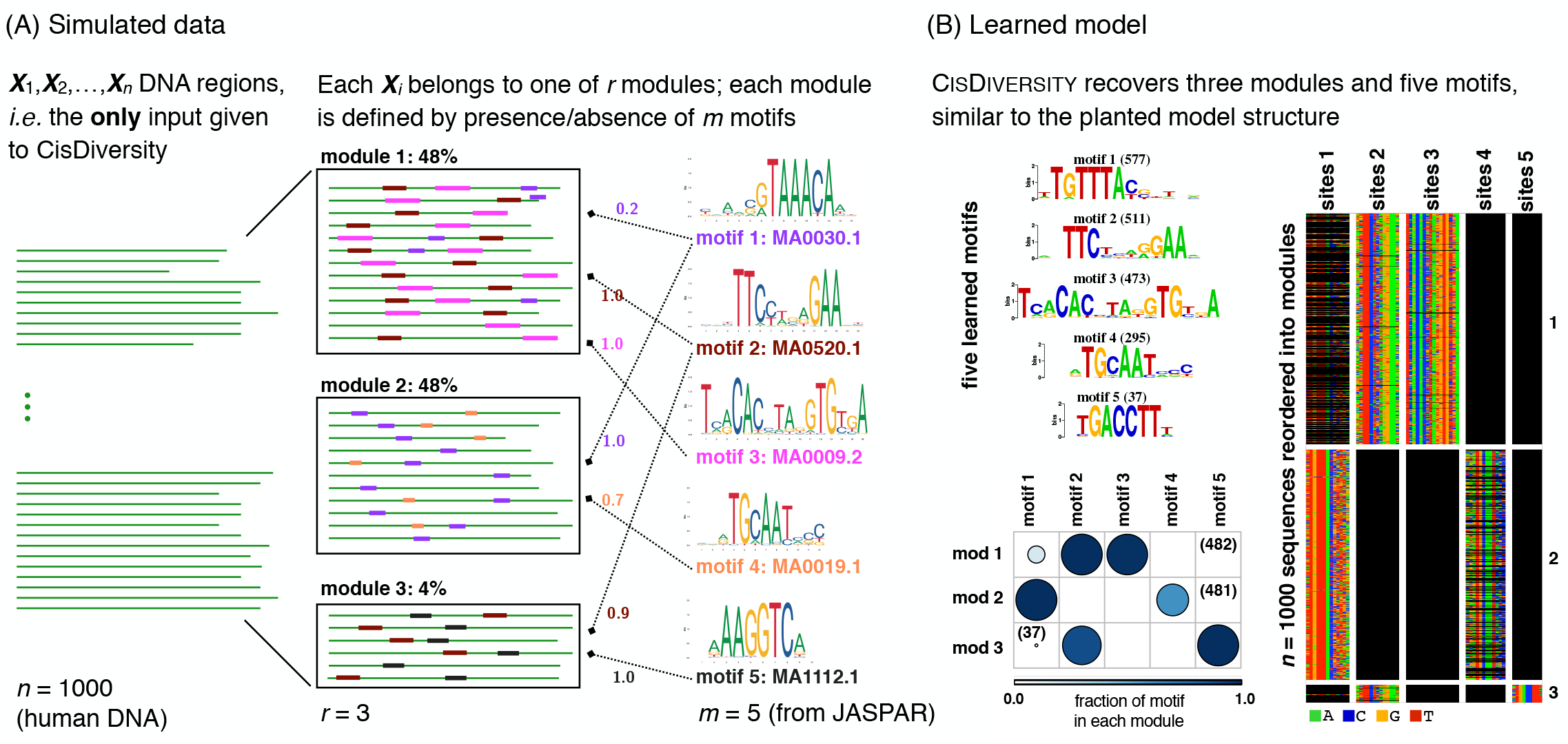
cisDiversity. (A) DNA regions reported by the experiment are given as input to cisDiversity. In this simulation, the *n* = 1000 regions are a mixture of three kinds of regions: each region resembles one of *r* = 3 regulatory modules. Each module can be represented in terms of the probability of occurrence of *m* = 5 motifs. For example, motif 1 is present in all sequences of module 2, 20% of sequences in module 1, but not at all in module 3. In contrast, motif 4 is present only in module 2, and that too only in 70% of the its sequences. (B) cisDiversity is run with upper bounds of *r* ≤ 10 and *m* ≤ 20. cisDiversity learns the planted structure in the dataset. The output has thr e components. First is the set of motifs that are learned, second (below) is *r* × *m* Bernoulli distributions describing the lear ed modules, while the third is an image matrix of the data, where each DNA sequence is a row, while the sites corresponding to each motif are represented in the column. If a site is absent, those cells in the column are shown in black. cisDiversity recovers the five motifs (motifs 1 and 3 are the reverse complements of the planted motifs) and the three modules to a great extent. The slight variablity in the number of sites and sequences in each module is expected due to the stochastic nature of both, the PWMs as well as the learning algorithm.

The algorithm behind cisDiversity uses Gibbs sampling to iteratively sample each of these unknown values with the aim of maximizing the posterior distribution (Methods). cisDiversity reports the output in three parts. The first is the set of de novo motifs ordered according to the number of sites that contribute to each motif. The second is the overall structure of the modules describing the contribution of each motif in every module. The color and the size of the circles denote the proportion of sites of the corresponding motif in each module. The last part of the output is the sequences clustered together as per the identified modules, displaying the sites that contribute to the PWMs in four colors for the four nucleotides. If a site is absent, those nucleotides are shown in black. The modules are ordered according to their size, the largest one shown on top. Figure 1B shows all three on the simulated dataset. cisDiversity finds all the motifs and the general module structure.

### 2.2 Performance on simulated datasets

cisDiversity can be thought of as a joint clustering and de novo motif discovery method. To systematically assess how well cisDiversity is able to retrieve modules and motifs, we simulated more such datasets. To better emulate reality, we used random non-repetitive regions from the human genome as the dataset ***X***. The number of planted motifs *m* was from the set {5, 10} and for each m, the number of modules was varied between the set {1, 2, 3, 5}. Now the performance of any clustering approach will depend on how separable the clusters are, while that of a motif discovery method will depend on how informative the motifs are. For the latter, we simply use real motifs from the JASPAR database (Methods). The former will be decided by the Bernoulli parameters (probability of presence of a motif) in each regulatory module. The more extreme (close to 0 or 1) this probability, the more informative the motif is in describing the module. A value closer to 0.5 for all *r* × *m* Bernoulli parameters will cause the modules to be less separable from each other. This variation was included in the simulated datasets by sampling the Bernoulli parameters from a beta distribution with a symmetric hyper-parameter *β* taken from the set {0.01, 0.1, 1, 10}. A value much smaller than 1 results in extreme values of Bernoulli probabilities, a value of 1 is akin to uniform sampling between 0 and 1, while a value of 10 results in probabilities closer to 0.5 and hence more “confused” modules. 10 sets were generated for each combination of parameters (*m*, *r*, *β*), resulting in 320 simulated sets, and cisDiversity was run on each. Module recovery was measured by the adjusted Rand index (ARI), popularly used to compare the similarity of two clusterings of the same data. Two identical clusterings get an ARI value of 1, while two random clusterings get a value of 0. Predictably, the ability of cisDiversity to identify the module structure goes down as the beta hyper-parameter goes up (Figure 2A). It is reassuring to see the modules are picked up although the *m* and *r* used in model learning (20 and 10, respectively) are larger than the true number of planted motifs and modules.

**Figure 2:**
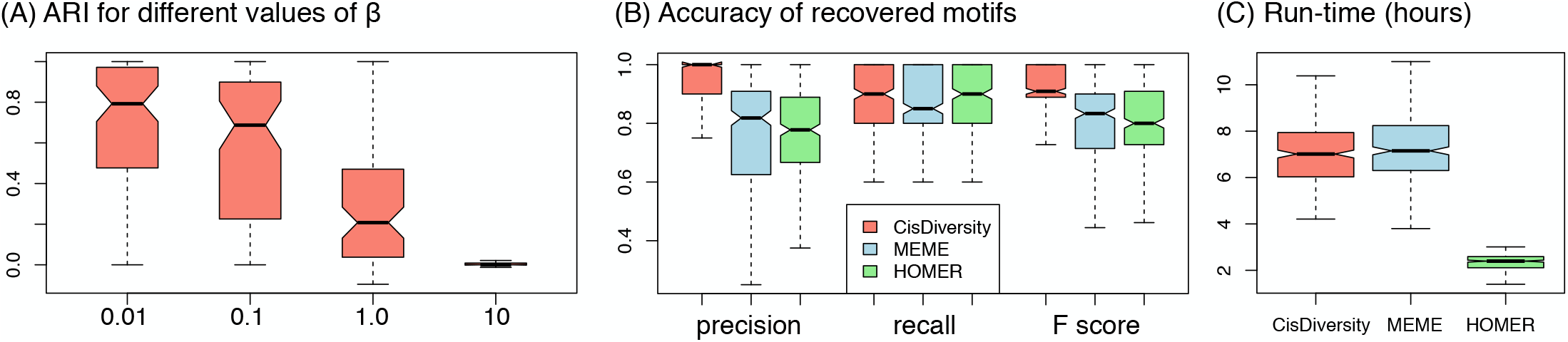
Performance on 320 simulated datasets. (A) Recovery of modules. Low values of *β* result into modules with more extreme (0 or 1) probability distributions of motifs. This is where cisDiversity does better in recovering the planted modules. For *β* = 10, the performance with respected to recovery of modules is similar to what a random clustering approach would do. (B) Recovery of motifs. Precision, recall, and F-score of recovered motifs across the 320 datasets for three different programs. (C) Time taken. All programs were run on a single core (Methods).

While plenty of methods solve the problem of de novo motif discovery, we are not aware of any other method that solves this problem of simultaneously identifying multiple modules. However, to evaluate how cisDiversity compares with state-of-the-art de novo motif discovery methods, we ran two commonly used programs—MEME^15^ and HOMER^16^—using default parameters on these datasets. The recovery of a planted motif was evaluated using TOMTOM ^17^, also from the MEME suite (Methods) in all cases. Figure 2B shows that cisDiversity performs better in terms of both precision and recall. However, we note that this is not a fair comparison, since the data does contain modules and no motif discovery method accounts for this fact. That said, when diverse modules is not a feature of the data, i.e., all datasets where the planted *r* = 1 or when the modules are close to being indistinguishable (*β* = 10), cisDiversity still is highly competitive (Figure S1). This shows that even if the data does not contain distinguishable modules, cisDiversity is capable of finding motifs. All programs were run in their serial mode, using a single core on an Intel Xeon CPU E5-2630 v3 machine. HOMER is by far the fastest, with there being not much difference between MEME and cisDiversity (Figure 2C).

In the next sections we apply cisDiversity to datasets arising from a range of different types of high-throughput assays. In each case we use a 200bp neighborhood of the summit or midpoint of each reported region, mask repeats and remove overlapping regions from the set (Methods). cisDiversity is applied with the same default options to all these sets, but with *m* = 30 and *r* = 15.

### 2.3 CTCF-bound regions display remarkable sequence level diversity

CTCF is a highly conserved TF, known to play different critical roles in regulation from binding insulators to forming chromatin loops in different contexts ^18,19^. Here we apply cisDiversity to see if these roles can be characterized in terms of modules from ChIP-seq data targeting CTCF. We used data from *D. melanogaster* in the white pre-pupa developmental stage ^20^. We also looked at the ChIP signal and the distance from the closest TSS at the sequences to see if the identified modules had specific properties/activities. Ni *et al.* ^20^ had identified a 9 bp motif `AGSKGGCGC` using MEME in the set, which resembles the canonical fly CTCF motif ^14^. This motif was present in approximately half of the regions. cisDiversity reports 17 motifs spread across 13 modules (Figure 3A–C). The top module—with most sequences—is dominated by the first motif, which matches the reported CTCF motif. This module also has a significantly higher (*p* < 10^−10^) ChIP signal, which is expected, if we assume this is where the binding of CTCF is strongest. Motif 1 contributes partially (27%) to module 4, where it co-occurs with motif 5. Motif 5 resembles the motif of another insulator binding protein suppressor of hairy wing, Su(Hw) ^14^. This module also has the maximum overlap with the ChIP-seq regions bound by Su(Hw) in the same developmental stage ^21^, suggesting that sequences in this module may be bound by Su(Hw) directly.

**Figure 3:**
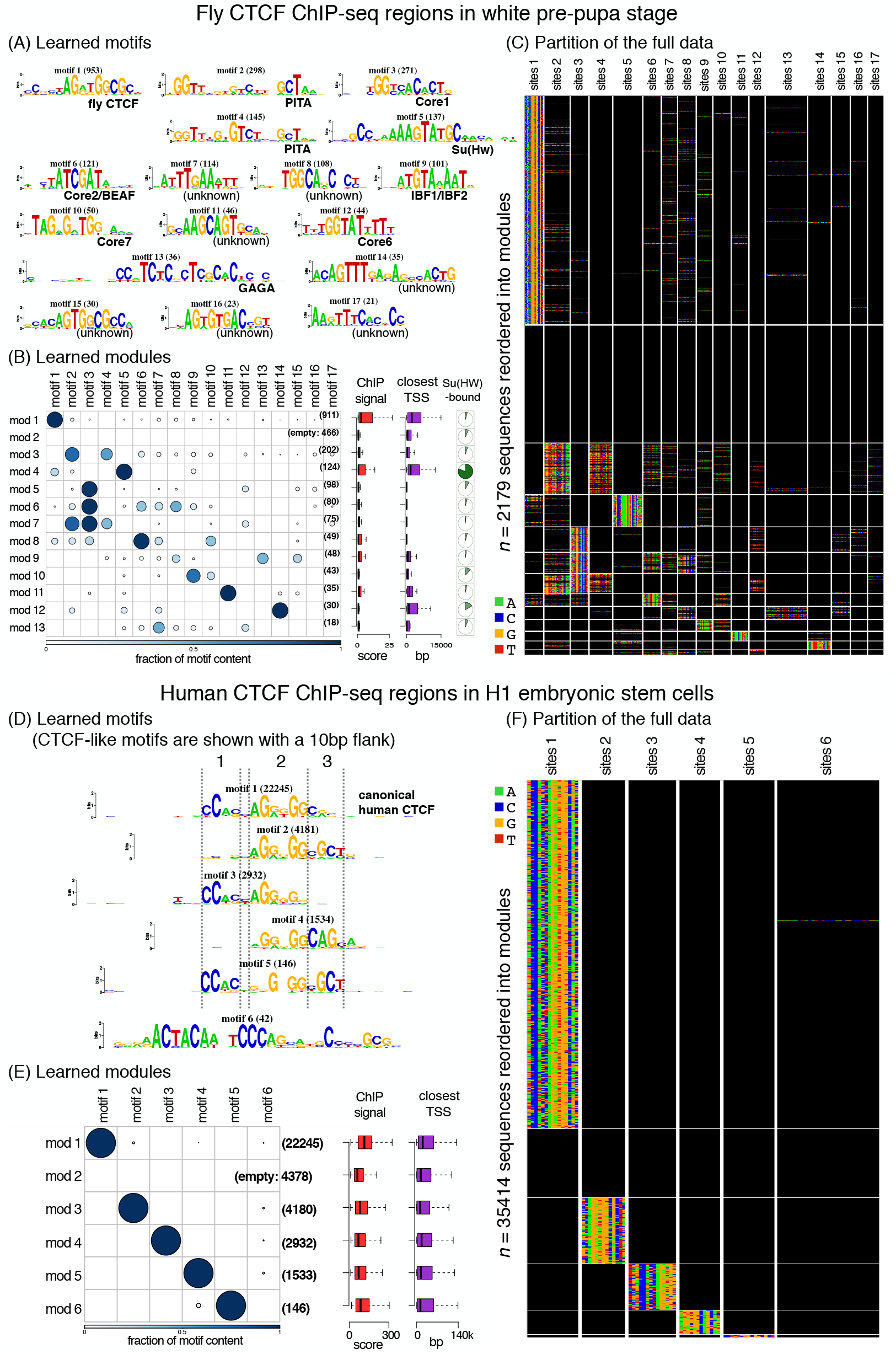
(A–C) cisDiversity identifies 17 motifs and 13 modules in the fly CTCF data. Motif 1 shown in a red box is the canonical fly CTCF motif. (D–E) cisDiversity identifies only seven motifs and six modules in the human CTCF data. Again, motif 1 in the red box matches the canonical vertebrate CTCF motif. Motifs 2–5 resemble the vertebrate CTCF, but differ at one of three parts denoted with dotted lines. These motifs are shown with 10bp flanks to ensure that they are genuine variants of the motif.

Multiple studies ^20,22^ have shown that in fly, the highest proportions of CTCF binding regions are in promoters. Indeed, if we look at this dataset as a whole, over half of the bound regions are within 1 kb of some TSS. However, when we look at the modules individually, we note that the two modules described above, which contain the CTCF motif in fact have fewer promoters (*p* < 10^−5^ for both). In contrast, modules 5–8 largely overlap promoters (< 100 bp from a TSS) and except for module 8 do not contain CTCF motifs. All of them are enriched with motif 3, a known core promoter element^23^, but with diverse co-factors. These are in fact different core-promoter architectures ^24,25^. Taken with the fact that ChIP signal is also low at these modules, CTCF probably makes indirect contacts here. Module 2 has no motif and has the lowest ChIP signal, which is also significantly lower than the other modules (*p* < 10^−5^), again suggesting non-specific or weak binding of CTCF.

Motifs 2 and 4 are highly similar and resemble the motif of the zinc finger TF PITA^26^. Both occur in modules 3 and 7. This implies that sequences in these modules have two copies of the PITA motif, and therefore both are required to describe the dataset. Now cisDiversity does not take into account the spatial distribution of the motifs in its model. However, in over two-thirds of the sequences the distance between the two copies is less than 50bp, suggesting cooperative binding.

Unlike vertebrates, there is no evidence to support cohesin-CTCF mediated chromatin loop formations in fly ^19^. Instead, it is believed that looping may be mediated by interactions of CTCF with other insulator binding proteins like Su(Hw), BEAF-32, IBF1/2, and GAGA^27^, all of which are identified by cisDiversity (motifs 5, 6, 9, and 13 in Figure 3A). We were unable to find strong matches for the other discovered motifs to any known TF motif in literature. However, sites at many of these motifs are conserved across flies (Figure S2), which suggests they may play a role in CTCF-related regulation.

These results are much in contrast with those obtained on human CTCF. cisDiversity was run on ≈ 35000 ENCODE CTCF ChIP-seq regions from human H1 embryonic stem cells (ESCs). Although the number of bound sequences is far greater than in fly, cisDiversity reports only six modules and six motifs (Figure 3D–F). The top motif is the canonical CTCF motif and four are variants with some parts of the motif missing or displaying nucleotide dependencies. These variants possibly correspond to the variable usage of CTCF zinc fingers ^18^; however, while nucleotide dependencies within vertebrate CTCF have been well-documented ^28,29^, we have not previously seen variants that are missing parts of the motif. Furthermore, these five motifs almost always occur in separate modules and together explain over 85% of the sequences. The only non-CTCF motif matches that of ZNF143, a transcriptional activator that binds at promoters, associated with CTCF^30^ and in loop formations ^31^.

Of the sequences that contain ZNF143, almost 90% also contain a CTCF variant, implying that interaction at these sequences with CTCF is not necessarily indirect/via ZNF143. Compared with fly CTCF, there is far more variability in the human CTCF motif itself and it does not appear to make as many indirect or non-specific interactions with DNA. However, the ChIP signal is significantly higher at the module with the canonical motif, suggesting that the modules with the variants are less strongly occupied by CTCF. CTCF is known to form loops and topological domains in mammals ^18^. It is possible that the various zinc fingers of CTCF interact at different chromosomal locations thereby facilitating loops between them, but more experiments would be required to definitively establish this.

### 2.4 GR binds to diverse regions after activation

We next applied cisDiversity on the human glucocorticoid receptor (GR), which binds to thousands of sites in response to exposure to the glucocorticoid (GC) hormone cortisol. GR is understood to bind primarily to DNase hypersensitive sites (DHSs) ^32^ and often with other pioneer factors ^33,34^. McDowell *et al.* ^35^ have probed binding of multiple factors including GR, before and after treating A549 cells with the synthetic GC dexamethasone (dex).

Figure 4A shows the results of applying cisDiversity on 6694 GR bound regions identified 1 hr after dex treatment. Six modules and five motifs are identified. The largest module contains motif 1 that matches the GR motif. Modules 2–5 are dominated by the other four motifs, which match motifs of TFs that are known to play a role in recruiting GR to its binding sites ^33,34^.

**Figure 4:**
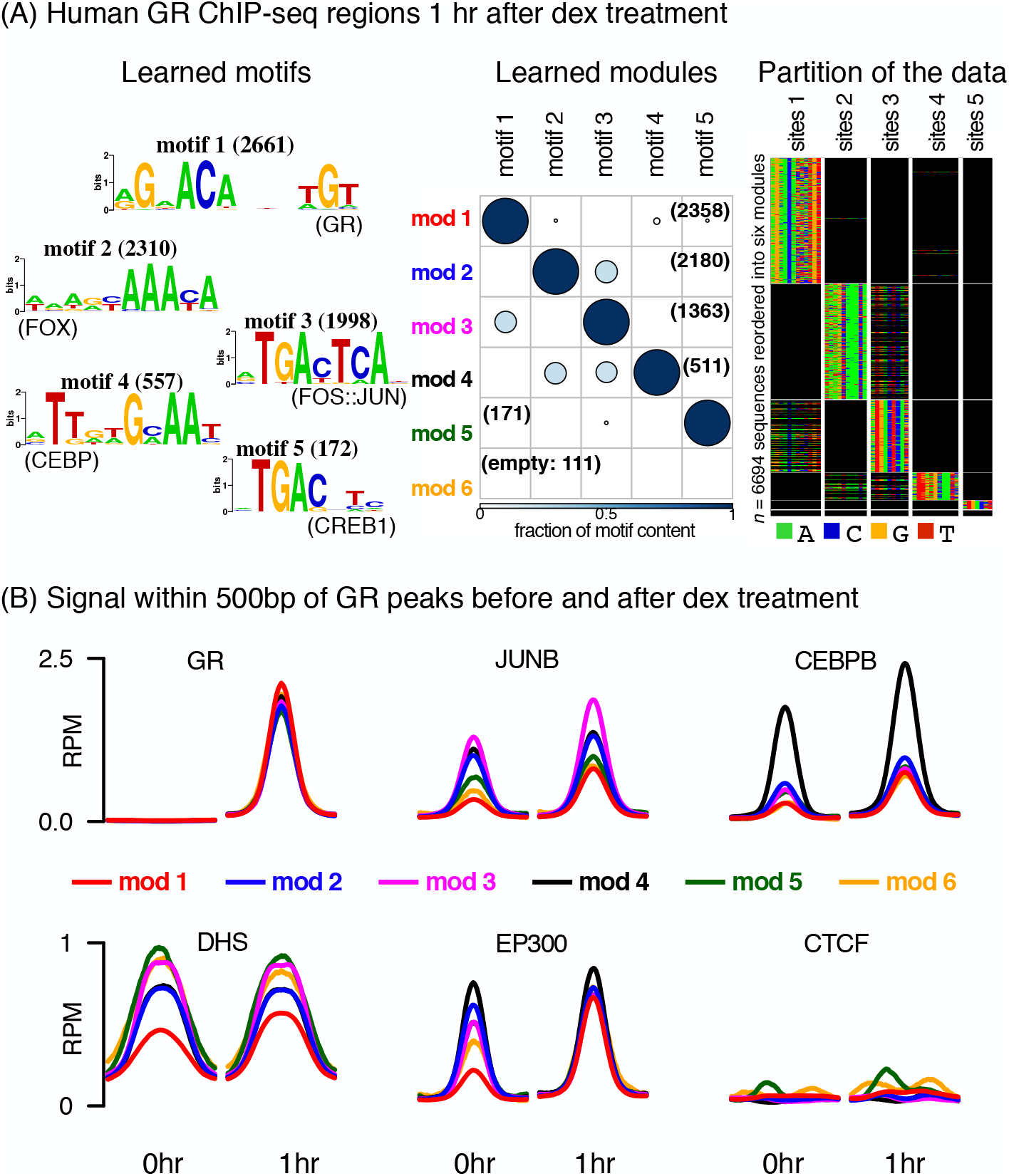
(A) cisDiversity identifies six modes and five motifs in the GR ChIP-seq regions. (B) The average DNase hypersensitivity signal and input-subtracted ChIP-seq signal in reads per million (RPM) of five TFs—GR, JUNB, CEBPB, EP300, and CTCF—before and after treatment at the GR-bound regions is shown for each module.

We looked at the ChIP-seq signal of profiled TFs at these modules both before and 1 hr after dex treatment. As expected, GR signal is non-existent before dex treatment at any module (no GR-bound regions were reported in this stage) and goes up at all modules after treatment (Figure 4B), but significantly more at module 1 (*p* < 10^−10^). Unsurprisingly, ChIP-seq signal of JUNB and CEBPB is higher at modules with their cognate sites. However, they appear to occupy those modules even before dex treatment, albeit with lower intensity. Their signal at all modules, including module 1 goes up after dex treatment, suggesting GR contributes to some sort of cooperative binding at all modules.

As has been noted before ^32^, all modules have some DNase hypersensitivity signal before treatment, but the difference before and after treatment is the most at the module with the GR motif. This difference is even more pronounced when we look at the enhancer mark: EP300 binding signal. McDowell *et al.* used the GR motif to scan the GR-bound regions and observed that regions with initial EP300 binding (before treatment) had a weaker median GR motif strength than did those without initial EP300 binding. Our results are consistent with this, but in addition they show that not only is the GR motif absent in regions with initial EP300 binding, but those regions can be explained with the help of other motifs. Interestingly, when cisDiversity is run on the EP300 ChIP-seq regions separately, it identifies near identical motifs, but with different module distributions; the module with GR motif is the fourth largest (Figure S3). CTCF ChIP-seq signal is plotted as a negative control: there is no difference before and after dex treatment of CTCF occupancy at these modules.

### 2.5 Enhancers have a different structure based on the detection assay

Enhancers play critical roles in activating gene transcription, while often being distant from the target gene. While certain TF binding sites are enriched at enhancers, there is no consensus sequence-based rule that can explain or characterize these regions. Over the last few years, several new high-throughput assays have been developed that measure a biochemical activity that is indicative of enhancers, either directly, or indirectly. For example, short, bidirectional, and largely unstable transcripts have been shown to originate at active enhancers ^36^. The function(s) of these enhancer RNAs or eRNAs are as yet not completely understood, but active TF binding at enhancers has been shown to increase corresponding eRNA levels ^37^. We applied cisDiversity to 14300 distal eRNAs ^38^ detected in the GRO-cap dataset from the K562 erythroid cell line ^39^. Figure 5A shows nine motifs and eight modules learned in this data. There are actually five more motifs, which together explain the ninth module of 32 sequences (Figure S4). The UCSC genome browser marks these sequences as segmental duplications ^40^. Indeed, clear conserved structures were identified by the multiple sequence alignment tool CLUSTAL^41^ when module 9 was given as input (Figure S5). For clarity, we have removed this module and the corresponding motifs here.

**Figure 5:**
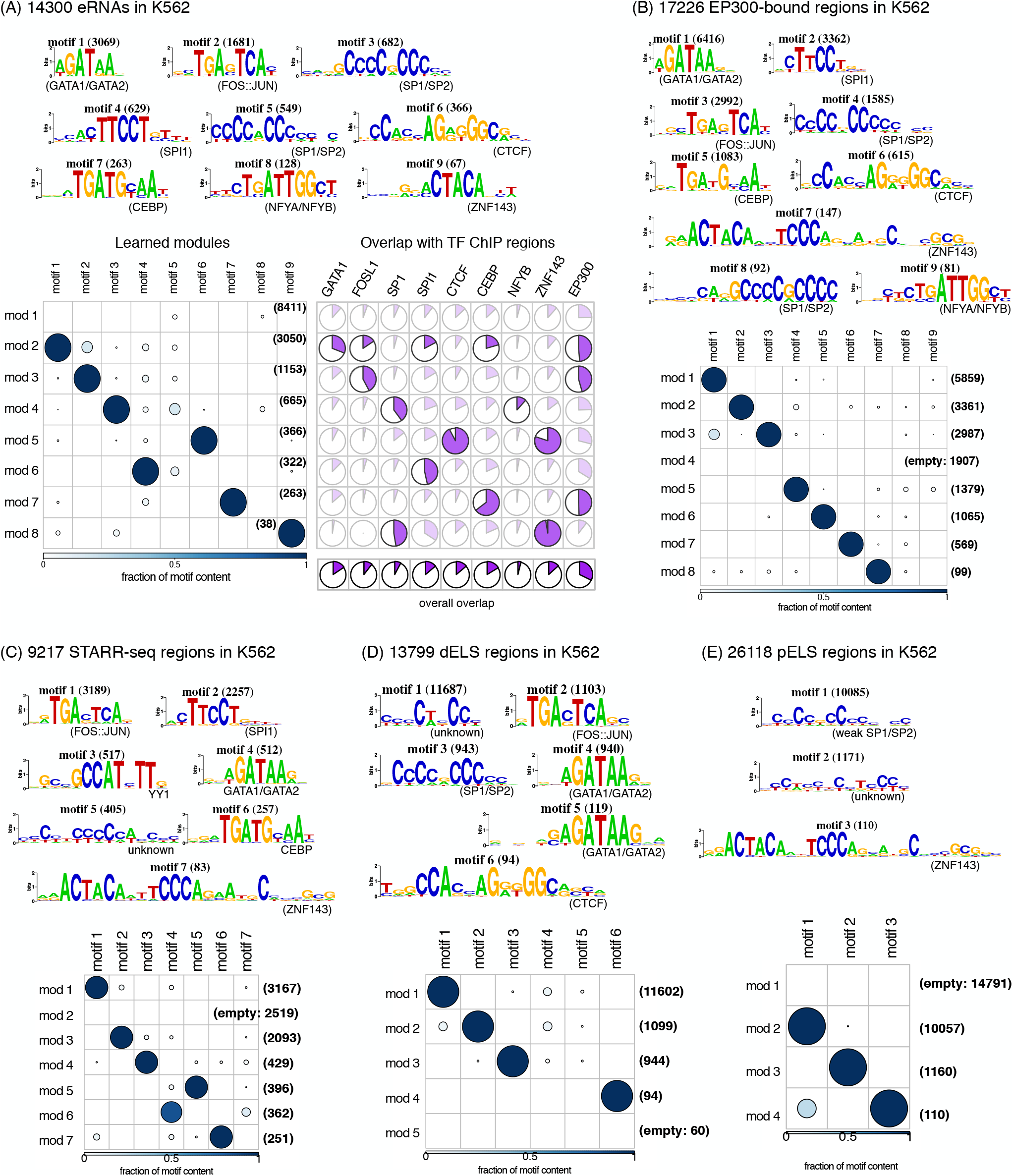
cisDiversity run on putative enhancers in K562. (A) On distant eRNAs, overlap with ChIP-seq data is significant (hypergeometric *p* < 10^−4^, shown in bold) in modules that contain the matching TF motif. Note that in some cases the overlap looks large but does not show up as significant, because the hypergeometric test corrects for the sizes of the overlaps and the modules. (B) Similar motifs are found in EP300 ChIP-seq data. (C) In STARR-seq peaks, YY1 is additionally discovered. CTCF and NFYA/NFYB are not enriched. (D) Distant and (E) Proximal enhancer-like sequences

In contrast to the models obtained from ChIP-seq regions, here the largest module, composed of over half the dataset, is largely empty, with no significant occurrences of any motifs. All other modules have one dominant motif, which matches a motif of a TF active in K562. We therefore assessed overlaps between modules and ChIP-seq regions of all these TFs. As expected, the overlap is significant (hypergeometric *p* < 10^−4^) for the corresponding TF, although the overall overlap with the complete set is, for most TFs, small. Overlap with the enhancer mark EP300 and active chromatin marks is higher (Figure S6), but there too we see differences in the degree of overlap across the modules. While EP300 mark is present at a large fraction of the eRNA sequences, it is significantly more at modules 2, 3, and 7, which are dominated by motifs of enhancer binding proteins GATA1, FOS::JUN (AP1), and CEBP, respectively.

The largely empty module in the distal eRNAs has no significant overlap with any of the external datasets. In fact, the overlaps are significantly *less* than that expected by chance (hypergeometric *p* < 10^−4^) with EP300 and other active chromatin marks. This, taken together with the fact that no significant sequence motif is detected in this module, suggests that perhaps these regions do not conform to the standard definition of enhancers.

Azofeifa *et al.* ^38^ had removed bidirectional transcripts that overlapped with annotated promoters to get the distal set of eRNAs. We applied cisDiversity to these 8324 excluded proximal transcripts and retrieved a completely different model with 10 modules and a much larger set of 25 motifs (Figure S7). Some motifs such as those matching SP1, NFYA/B, ZNF143 are common between both sets, but proximal eRNAs are enriched with many other promoter motifs such as YY1, RFX, CREB1, NRF1, etc. Contrary to the distal modules, there are more motifs per module, implying more cooperative binding at these regions, which is a hallmark of promoters ^42^. The enrichment of active chromatin marks is also different across these modules. These regions are more often accessible and have more overlaps with all active marks, except for H3K4me1, which is associated with enhancers ^43^ and less so with promoters ^44^. This is consistent with the dataset being separated based on overlaps with annotated promoters. However, even within the modules in both datasets, there are differences across the marks, implying that not all eRNAs have the same chromatin signatures, even after separating them as distal and proximal. But more importantly, these differences can be partially explained with modules and motifs.

We next ran cisDiversity on four other datasets, which also report enhancer activity in K562. The first is the EP300-bound sequences in K562, where we again get nine motifs, almost identical to ones in distal eRNAs, but present in different fractions of the sequences (Figure 5B). In contrast to the distal eRNA set, but similar to the other TF ChIP-seq datasets, a large majority of sequences contain some motif in the EP300 set. We next looked at STARR-seq data. In a STARR-seq assay, random genomic fragments are placed in the 3’ UTR of a reporter gene with a minimal promoter and the resulting plasmids are transfected into the cells of interest, K562 here. The enhancer activity of these fragments is then measured by sequencing the 3’ UTR of the reporter gene transcripts. Unlike the other methods, this assay considers regions outside of their chromatin context so it can report regions which are inaccessible, but have a potential for enhancer activity ^45^. We used the peaks reported by Lee *et al.* ^46^, using their STARRPeaker method, which resulted in a little over 9000 sequences. Interestingly, although there are very few sequences common between the STARR-seq regions and the eRNAs or EP300-bound regions (Figure S8), many motifs are common between the three sets. However, YY1 is discovered additionally in the STARR-seq regions (module 4), while NFYA/NFYB and CTCF are absent. Since STARR-seq assays considers regions outside of chromatin context, we looked at whether the modules had accessibility profiles distinct from the first two datasets. Indeed, modules 2 (devoid of motifs) and 4 (dominated by YY1) are significantly (*p* < 10^−5^) depleted of DHSs (not shown). This suggests that these modules are suppressed by endogenous chromatin in K562. On the other hand modules 1 (AP1-dominated) and 6 (GATA-dominated) are significantly enriched with DHSs: these structures were also identified in the eRNA and EP300 assays.

The last two datasets are from ENCODE phase III, where Moore *et al.* ^47^ have published a registry of candidate cis-regulatory elements based on results from multiple high-throughput experiments. They report two disjoint subsets, which they propose have distal enhancer-like signatures (dELSs) and proximal enhancer-like signatures (pELSs). The criteria for a sequence to be included in either of these sets is that it should have high DNase and H3K27ac signals. pELSs are within 2000bp of an annotated TSS, while dELSs are away. pELSs additionally must have low relative H3K4me3 signal, to ensure they are not active promoters. Other than an SP1/SP2 motif, there are no common motifs in the two sets (Figure 5C,D). CTCF is found in the dELS while ZNF143 in pELS, which is not surprising since these TFs’ binding sites are enriched in regions distal and proximal to TSSs, respectively ^48,49^. In contrast to the first three enhancer datasets, cisDiversity finds no motifs matching SPI1, CEBP or NFYA/B. No GATA motif is found in the pELS set, possibly because GATA proteins primarily bind to distal enhancers ^50^. Overall, fewer signatures that look like TF-motifs are identified in these enhancer-like regions from ENCODE III.

### 2.6 Modules in open regions have differing future fates

We next looked at data from two different technologies that measure chromatin accessibility. The first is the assay for transposase accessible chromatin with sequencing (ATAC-seq) and the second is the one that measures DNase accessibility (DNase-seq). Cusanovich *et al.* ^51^ used single-cell ATAC-seq on *Drosophila* embryos at three different stages after egg laying. Here we report cisDiversity results on the earliest stage: 2–4hrs after egg laying. We considered regions that were open in at least 10% of the cells assayed at that stage. This resulted in 9963 unique peaks. cisDiversity finds 28 motifs and 13 modules (Figure S9). For clarity, Figure 6A shows only those motifs that contribute to at least a quarter of the sequences in some module. Without any additional information, cisDiversity largely partitions the data into modules that are significantly enriched with promoters (within 500bp of an annotated gene) and those that are significantly depleted (*p* < 10^−4^). Most of the motifs that contribute to the promoter-enriched modules are well-established core-promoter motifs. The two PITA motifs from fly CTCF data (Figure 3A,B) are recovered here as well, in module 10. Modules distant from TSSs are potentially enhancers and/or insulators. Indeed, enrichment of CTCF and Su(Hw) in module 11 suggests this module harbors insulators. Similarly, dinucleotide repeats of GA (bound by GAGA/TRL) and CA are known to be features of fly enhancers ^52^. One of the promoter modules comprises of tRNAs and cisDiversity captures the highly conserved hair-loop structure of tRNAs as a “motif” (module 12).

**Figure 6:**
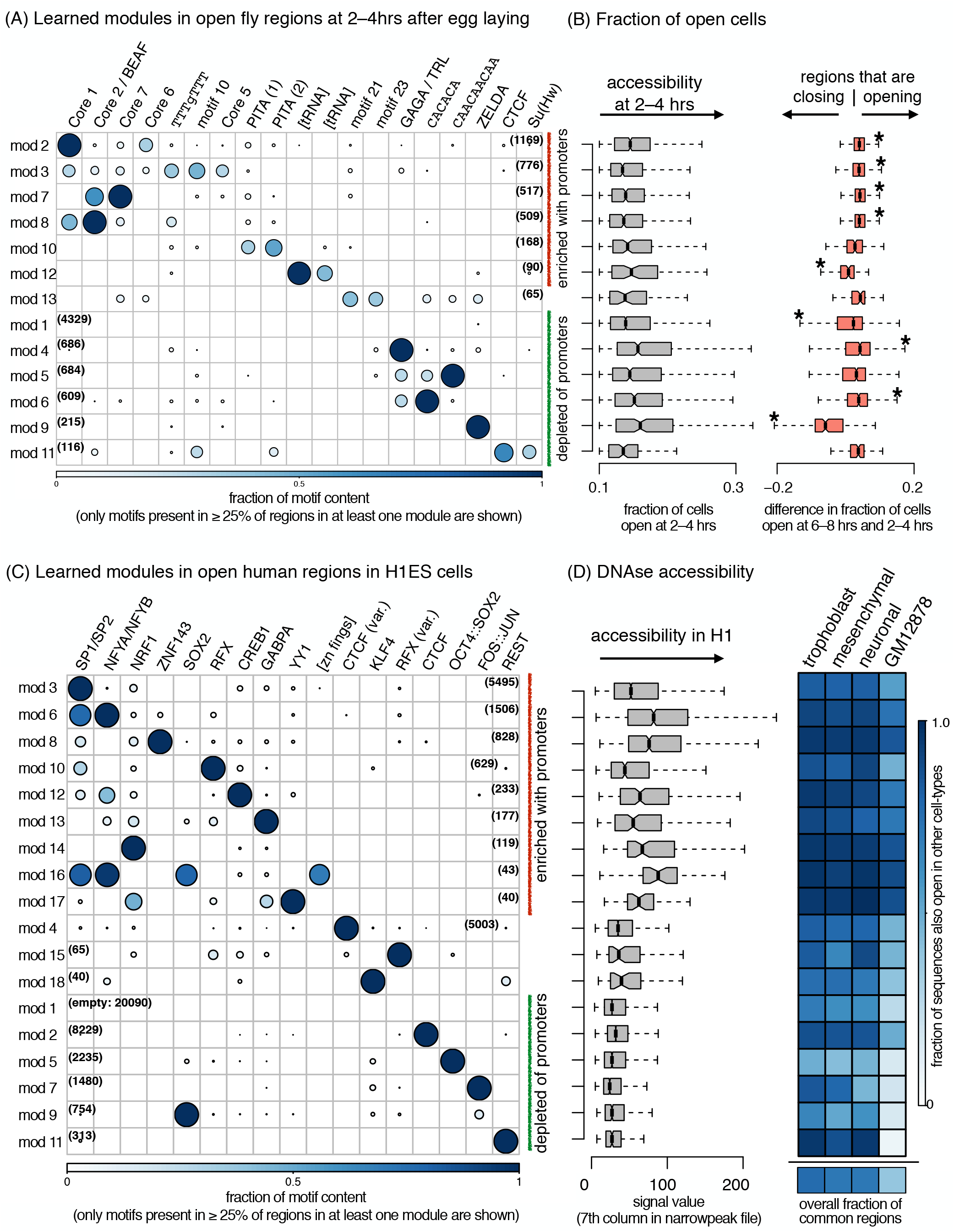
cisDiversity is run on open regions. (A) 13 modules and 28 motifs (Figure S9) are learned on ATAC-seq regions, which are open in at least 10% of the cells probed 2–4hrs after egg laying. Only the 19 motifs that contribute to at least a quarter of the sequences in some module are shown here for clarity. Modules are reordered: the red and green modules are significantly (hypergeometric *p* < 10^−4^) enriched with promoters and depleted of them, respectively. (B) Grey: there are a few differences in the fraction of cells open within each module. Orange: modules 2, 3, 7, 8, 4, and 6 are significantly more open in the cells 6–8 hrs after egg-laying, while modules 12, 1, and 9 are closing at that time point. (C) 18 modules and 21 motifs (Figure S10) are learned on DNase-seq regions in H1 ESCs. Again, only the motifs appearing in a quarter of sequences of some module are shown here. Red and green modules are as in A. (D) Grey: promoter modules have a higher DNase signal in general, but there are variations among them. Blue: fraction of each module (and total below) that is also open in trophoblast, mesenchymal, and neuronal stem cells (all derived from H1 ESCs) and GM12878 shows considerable variation across modules and cell-types.

There is little difference in the fraction of cells open at the regions across the modules: only modules 4 and 6 (distal regions with dinucleotide repeats) are significantly more open. However, when we compare the fraction of cells open at these regions with the fraction open four hours later (6–8hrs after egg laying), we see several modules significantly changing their accessibility (Figure 6B). Almost all promoter modules are opening up, except for the tRNAs. We found no evidence of tRNA gene expression going down during these stages of development. However tRNA genes are known to play a role in remodeling chromatin, and act as insulators or harbor origins of replication in other organisms ^53,54^. Modules 1 and 9 are the only other modules that are significantly closing. Module 1 has very few predicted binding sites, with no motif that occurs in even 10% of the sequences. This is in concordance with recent results that suggest two classes of enhancers are active during early *Drosophila* embryogenesis, one of which has significanlty lower TF occupancy, in general ^55^. All sequences in module 9 however, contain a motif matching that of Zelda. Zelda is a pioneer TF that is critical for the maternal to zygotic transition, which happens during this stage ^56^. This suggests that these Zelda-enriched regions are less accessible in the subsequent stages.

Figure 6C shows the modules learned on 47279 DNase hypersensitive regions of H1 ESCs. The original option of *r* = 15 resulted in 15 modules, so cisDiversity was re-run with *r* = 20, reporting a total of 18 modules. As in fly data, here too, the modules are significantly enriched with promoters or depleted of them. All promoter modules have high DNase accessibility, but there are significant differences between them. For example, while modules 3 and 6, both are enriched with promoters and with the SP1/SP2 motif, module 6, which also contains an NFYA/NFYB motif, has a significantly (*p* < 10^−10^) higher DNase hypersensitivity signal. Module 16 is primarily composed of core-promoters of a KRAB family of zinc fingers, with a distinct long motif. Modules depleted of promoters are typically characterized with motifs of SOX2, OCT4::SOX2, which are H1 ESC-specific TFs and motifs of other pioneer factors AP1, REST, and CTCF.

We looked at the DNase accessibility at all these regions in stem cells derived from H1 ESCs available in ENCODE: trophoblast, mesenchymal, and neuronal stem cells (Figure 6D, Figure S10). While the overall fraction of regions overlapping with DHSs in these stem cells is similar (0.76 for trophoblast, 0.71 for mesenchymal and neuronal), the overlap-fraction within modules ranges from 0.42 to 1.0. Indeed, over 90% of sequences of modules 8, 12, 14, 16, and 17 are accessible in these three stem cells. In contrast, modules dominated by motifs of pioneering ES TFs (SOX2, OCT4, and KLF4) are less accessible, which is expected. Module 11, containing the REST motif, has a high accessibility in all the stem cells. The canonical CTCF motif (module 2) is significantly more often accessible than the variant (module 4) in the other cells, although in H1 ESCs, the variant has a higher accessibility signal (*p* < 10^−5^). DHSs in the lymphoblastoid GM12878 cell-line (derived from blood) are used as control. Overall, the overlap is lower for all modules, but we see the same trend of higher overlaps at promoters and lesser ones at distant modules.

## 3 Discussion

The importance of combinatorial binding of TFs in gene regulation is well-established ^57^ and several attempts have been made to detect regulatory modules from high-throughput data. One of the earliest motif-module detection programs was CisModule ^58^, where the goal was to learn the location of a module in each sequence along with motifs. The module, however, was of a single kind. Self organizing maps ^59^, non-negative matrix factorization ^60^, as well as topic models ^61^ have been subsequently used to cluster regions based on multiple ChIP-seq datasets, with the goal of identifying different modules. These methods have been used to predict complexes forming along the chromatin and co-localization of TFs. However, they explicitly require regulatory information in terms of other high-throughput experiments for each region, which act as features for clustering. They do not incorporate sequence information or motifs as part of the model. cisDiversity is the first attempt to cluster regions based on motifs that are themselves learned during the process, requiring no additional experimental or TF binding information. The discovered modules indeed correlate with experimental binding data of TFs with matching motifs, showing that regions contain sequence information that can characterize functional modules. On simulated sets, as a motif discovery method, it performs better than standard approaches especially in terms of precision. We believe this is because of its model-based approach, where the goal is to learn a model that explains the full dataset, and not learn motifs that individually explain a portion of the sequences.

Regulatory modules are typically studied in accessible regions or putative enhancers, assuming cooperativity of TFs there. However, cisDiversity gives new insights even in the ChIP-seq datasets investigated here. Take for example the heavily studied CTCF protein. The original fly study ^20^ also mentioned that many sequences did not contribute to the overrepresented CTCF motif, but all subsequent location and evolution analysis was done considering all ChIP regions equally. However, cisDiversity clearly shows that some sequences are core-promoters with no CTCF motif, while some are enriched with other insulator-binding TF motifs. The evolutionary profile of the motifs is also diverse in the dataset. On the other hand, the human CTCF behaves remarkably differently, with ZNF143 being the only non-CTCF motif that is discovered. Instead, several individual variants of the CTCF motif are enriched in different modules, suggesting a possibility of differential usage of its zinc fingers while binding DNA.

Putative enhancers have been detected using multiple high-throughput assays in the same cell-type or context. The datasets arising from these assays differ in terms of their cardinality, length distributions, and evolutionary features ^62^. cisDiversity shows the differences in terms of motifs and modules. No motif is common across all the enhancer-detection strategies, at least in the cell-type assessed here. It is important to note that each dataset used here comes from an assay that measures a different biochemical activity or combination of activities. cisDiversity can be used to further tease out the differences between the sequences reported by these strategies in terms of motifs and their combinations.

cisDiversity automatically clusters accessible regions into promoters and distant regions. This is not surprising since the sequence architecture of promoters is different. However, even within promoters and putative insulators/enhancers it captures considerable sequence-level diversity. While the modules in Figure 6 are reordered based on their propensity to have more or less promoters, no module is completely composed of or devoid of promoters. Distant sequences in a promoter-enriched module should be further studied for potential promoter activity and vice versa.

The model does not incorporate any information about known motifs, their multiplicity or distance between them. These aspects can be studied from the learned parameters. For example, although not modeled explicitly, cisDiversity identifies homo-typic binding if it appears in a significant fraction of sequences, by learning more than one copy of the same motif. The pair of PITA motifs (Figures 3A and 6A) has not been reported before. The distance between motifs and relative orientation can be assessed as well. Similarly, the distance of each motif from the summit of the peak can give insights into the likelihood of direct binding of the profiled TF at that motif ^63^. Although not shown here, cisDiversity can be restricted to look for stranded motifs especially when dealing with strand-specific datasets like 5’ CAGE TSSs or UTRs. In all results described here, we have reported the top scoring model, treating it as a hard clustering method. However, we note that cisDiversity learns a probability distribution over the modules for each sequence and multiple module usage can also be explored ^61^. Currently cisDiversity reports no significance value or false discovery rate for a learned model. However, it does report the posterior probability associated with the learned model. These values are far lower in datasets with random regions with no planted modules/modules than those used in Section 2.2 (Figure S11). Such distributions can, in principle, be learned to report an associated empirical “p-value”.

The strength of cisDiversity is in its lack of reliance on known PWMs or modules, making it general enough to be used on any set of DNA sequences. It is the principle of treating data—even when it is from a single high-throughput experiment—as a diverse set, but at the same time trying to explain as much of it as possible, using sequence signatures, that makes cisDiversity unique.

## 4 Materials and Methods

### 4.1 Model framework

The goal of most high-throughput experiments is to identify all regulatory regions with a certain biochemical property in a specific context. These regions might display that property due to *r* different mechanisms and we assume one of these mechanisms is encoded in the DNA sequence of each region. A “mechanism” is represented as what is commonly known as a regulatory module. Each of the *r* modules is characterized by the presence or absence of *m* motifs. Each level of the model *M*, characterized by *r* and *m* is described below, in a bottom-up manner:

1. Motif level We assume at most *m* motifs are present and contribute to modules, across the complete set of reported regions. Motif *k* has a width of *w*_*k*_ and is characterized as a product of w_*k*_ categorical distributions over the four bases, as is commonly done in a position weight matrix. ***ϕ***_*k*_ is used to denote these distributions, where 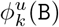 is the probability of finding base `B` at position *u* within motif *k*, `B ∈ {A, C, G, T}`. Given any *w*_*k*_ length DNA sequence 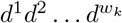, the likelihood of it being an instance of motif *k* is 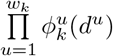.
2. Module level We assume at most *r* modules are present across the complete set of reported regions. Each module is modeled as a product of Bernoulli distributions over the *m* motifs: module j has a probability *f*_*jk*_ of containing motif *k* and a probability (1 − *f*_*jk*_) of not containing it.
3. Region level The experiment reports a total of n genomic regions ***X***_1_, ***X***_2_ …, ***X***_*n*_. Region ***X***_*i*_ is a DNA sequence of length *l*_*i*_: 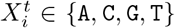, where 1 ≤ *t* ≤ *l*_*i*_. Parts of each sequence are instances of a subset of the *m* motifs; others parts, i.e. the “background”, are modelled using a second order Markov chain over the nucleotides, whose parameters are denoted by ***ϕ***_0_. *Z*_*ik*_ denotes the position within ***X***_*i*_, where motif *k* is present; i.e. 1 ≤ *Z*_*ik*_ ≤ *l*_*i*_ - *w*_*k*_ + 1. *Z*_*ik*_ = −1 in cases where there is no motif *k* in sequence ***X***_*i*_. Motif occurrences are not allowed to overlap and each motif can have at most one occurrence in each sequence. Sequence ***X***_*i*_ has a module identity *I*_*i*_ that is modeled with a categorical distribution *γ* over the *r* modules: *γ*_*j*_ is the probability of a sequence having a module identity *j*.

The set of regions ***X*** is all we are given, which we use first to compute the background Markov model as well as the lengths ***l***. We are also told whether motifs can occur on either strand or on the given strand only. The default is the former, in which case the sequences are appended to their reverse complements and their lengths are effectively doubled. We assume we know the structure of the model, i.e. *r* and *m* are given. The unknown parameters **Θ** are therefore *ϕ*_1…*m*_, ***w***, ***Z***, ***f***, ***γ***, and ***I***. The likelihood of region ***X***_*i*_, based on these assumptions can be computed as:

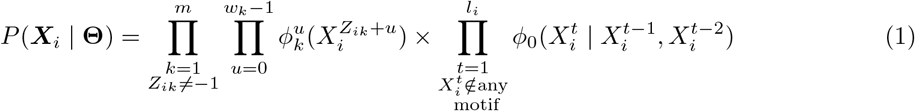

The first term denotes the probability associated with all the motifs that are present in ***X***_*i*_, while the other explains the sequence not overlapping with any of the motifs, using the backround Markov model. cisDiversity uses the second order Markov model as default, where each sequence has a background model built only from its 3-mers, to accommodate the vast heterogeneity in eukaryotic sequences^28^. But this can be changed by the user if required.

The complete likelihood is simply:

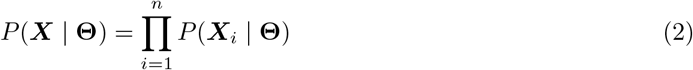

Given a set ***X***, the goal is to find the **Θ** that maximizes the posterior distribution, which can be computed as:

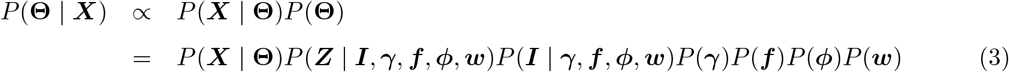

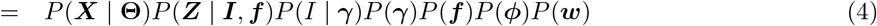

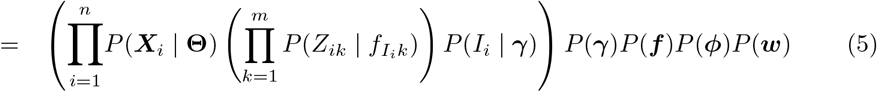

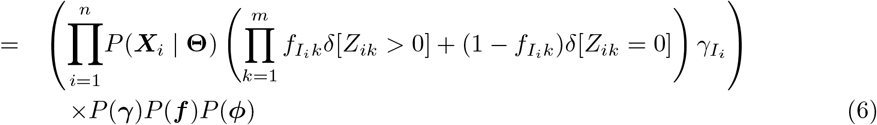

where *δ*[*condition*] = 1 only when *condition* is satisfied and 0 otherwise. The simplification in equation 4 arises due to the structure of the model: the site positions ***Z*** are independent of ***γ***, ***ϕ***, ***w*** when conditional on the module identity ***I*** and its associated probabilities ***f***; ***I*** is independent of ***f***, ***ϕ***, ***w***, when its categorical distribution ***γ*** is known. The second term in equation (5) is the product of Bernoulli probabilities, which are assumed to be independent. The third term is simply the categorical probability of the module identity. The rest are independent priors over ***γ***, ***f***, and ***ϕ***, which are discussed in the next section. The prior over the widths is considered uniform.

### 4.2 Model learning

In order to find the value of **Θ** that maximizes equation (6), we need to design appropriate priors over the parameters. We assume conjugate symmetric Dirichlet priors over all categorical parameters: (1) prior over ***γ***, which characterizes the distribution of module identity ***I***, has *r* equal hyperparameters defined by *α*_module_ (1 by default); (2) prior over each *f*_*jk*_, which characterizes the distribution of presence or absence of motif *k* in module *j*, has hyperparameters, defined by *α*_YES_ and *α*_NO_ (both set 0.1 by default); and (3) prior over each 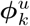, which characterizes the distribution over the four nucleotides at position *u* in motif *k*, has four equal hyperparameters defined by *α*_PWM_ for all *u* and *k* (all set to 0.1 by default). Hyper-parameters less than 1 assume extreme final distributions: we expect motifs to be informative, i.e. we expect nucleotide probabilities to be close to 0 or 1. Similarly we expect modules to be specifically described with the presence or absence of motifs. However, we do not know the number of non-empty modules, in other words, the inherent diversity in the data. Therefore we set *α*_module_ to 1 by default, which assumes all distributions are equally likely a priori. However, all these values can be changed by the user.

We use Gibbs sampling to draw samples from the posterior distribution with the aim of learning the parameter values that maximize it. In order to reduce the search space and speed up the sampling, we use collapsed Gibbs sampling^64^, marginalizing over ***f***, ***ϕ***, and ***γ***, while sampling only the other unknowns. The objective function then reduces to:

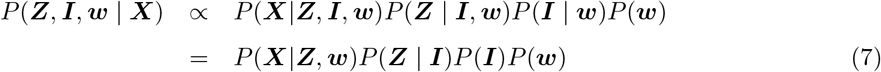

We therefore need to iteratively sample *Z*_*ik*_, *I*_*i*_, *w*_*k*_ for each *i* and *k*. If we assume a binding site is equally likely to be anywhere within a sequence, after dividing the posterior distribution by the background for each sequence, we can compute the sampling expressions for *Z*_*ik*_ as:

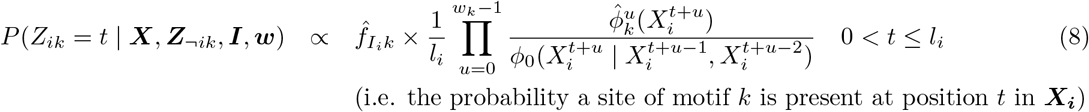

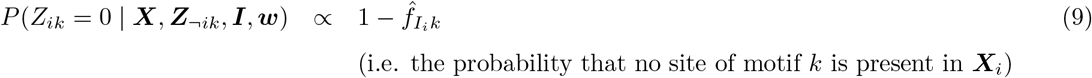

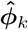 and 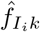 are the posterior probabilities after marginalizing, i.e. integrating out ***ϕ***_*k*_ and 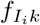 from equation (7) when sampling *P*(*Z*_*ik*_ | ***X***, **Θ** \ *Z*_*ik*_), assuming Dirichlet priors over both:

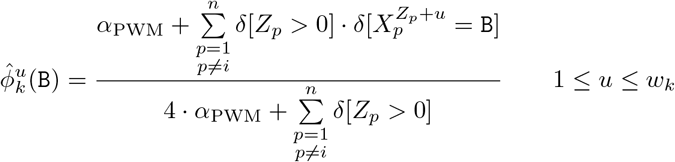

and similarly:

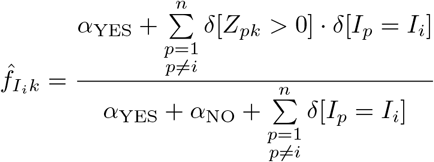

We note that the first term of equation (8) is similar to standard collapsed Gibbs sampling applied to motif discovery, except that the counts are computed only from those ***X***_*p*_ that contain a motif *k* at the current iteration, while the second terms arises from the contribution motif *k* makes to the current module *I*_*i*_.

The sampling expression for *I*_*i*_ can be similarly derived by collapsing ***γ*** and ***f*** from equation (7):

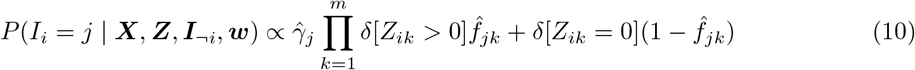

where 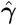 is the posterior probability distribution after integrating out ***γ***:

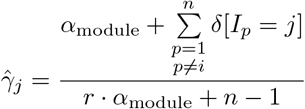

The width *w*_*k*_ of motif *k* is sampled a little differently. Instead of looking at all possible widths at one time, the width is allowed to increase or decrease by one on either side or stay the same as done before ^65^.

Each of *Z*_*ik*_, *I*_*i*_, and *w*_*k*_ are sampled iteratively using the expressions as described. The full posterior probability (equation 7) is computed after each sampling step and stored if found to be the best thus far. If the probability does not increase for a predetermined number of iterations (500 by default), the sampling stops and a hill climbing approach ^65^ starting from the sample with highest probability is taken until the probability stops increasing. The final model is cleaned up: all motifs that are composed of less than a minimum number of sites (20 by default, can be set to 0 by the user) are emptied out and all modules with less than minimum number of sequences (20 by default, can be set to 0 by the user) are clubbbed together. By default, cisDiversity starts from 10 different initializations and reports the model that scores the best across the 10 trials.

### 4.3 Simulated datasets

A total of 320 datasets were simulated for testing the efficacy of cisDiversity. The JASPAR2018 CORE vertebrate motifs were used for this purpose, regardless of their information content. However, many of these motifs are similar to each other. To create a non-redundant set, going serially, we removed all motifs that had a match with any earlier one according to the motif comparison tool TOMTOM ^17^. This resulted in a total of 189 JASPAR motifs. Each simulated dataset had 1000 DNA sequences of length 200 bp each, sampled randomly from the non-repetitive part of the human genome. Each dataset was constructed as an instance of the model described earlier. At the motif level, *m* motifs were drawn at random from the set of non-redundant motifs. At the module level, the Bernoulli parameters *f*_*jk*_ were sampled from Beta distributions with two symmetric (equal) parameters (*α*_YES_, *α*_NO_). At the sequence level, the categorical distribution *γ* was first sampled using the uniform parameter value of *α*_module_ = 1 and set for that model instance. For each sequence, this *γ* was used to first sample the module *j*. Then, the *f*_*jk*_ were used to sample whether motif *k* was to be included in the sequence or not. If yes, a site was sampled from the PWM *k* and randomly planted in the sequence, on either strand with equal chance, ensuring no overlap with any other planted site. Multiple datasets were created: *m* was one of {5,10}, *r* was one of {1,2,3,5}, *α*_YES_ was one of {0.01,0.1,1.0,10.0}. There were 10 dataset instances of each parameter set of {*m*, *r*, *α*_YES_} resulting in a total of 320 different datasets on which cisDiversity and other programs were tested.

We note that two modules can have very similar Bernoulli distributions, making them less separable, but the models were not screened for this possibility because (a) that would be rare with *m* ≥ 5 and (b) this would only mean we are underestimating cisDiversity’s performance.

cisDiversity was run with *m* = 20 and *r* = 10. MEME^15^ was run on these datasets using the command line options: `-nmotifs 20 -maxsize 100000000 -revcomp -dna -evt 10`. MEME was run with background Markov models of orders zero through five separately, learned on the non-repetitive human DNA sequences used to create the datasets. Markov order of three gave the best F-scores; those results are reported here. HOMER^16^ was run using the command line options: `-noknown -nogo -basic`. HOMER was also given the background file of the non-repetitive human DNA sequences. TOMTOM was used to compare the learned motifs with the JASPAR planted motifs. An E-value of 0.01 was considered a match.

### 4.4 Biological datasets

All datasets were processed in the following manner. 200 bp neighborhood of the summits (if reported, else the midpoint) of the regions were extracted. If two regions overlapped, only the first one in the list was kept. This was done to ensure that the model would not identify the same site in genome twice. Only those regions with at least 100 bp of non-repetitive bases in them were retained and used as input to cisDiversity. All datasets with their accession numbers are listed in Table S1. cisDiversity was run on each dataset with *m* = 30 and *r* = 15. All other parameters were set to default values. Reported modules were assessed for enrichment with promoters, assay signal, etc with either a Wilcoxon test or a hypergeometric test. All p-values in the text have been reported after Bonferroni multiple hypothesis correction. Sequence conservation plots were computed from phastCons scores ^40^.

## 5 Acknowledgments

We thank Rahul Siddharthan for useful discussions. This study was supported by grants from DBT Government of India BT/PR16240/BID/7/575/2016 and BT/IN/BMBF-BioHr/32/LN/2018-19.

## References

[1] Johnson, D. S., Mortazavi, A., Myers, R. M. & Wold, B. Genome-wide mapping of in vivo protein-DNA interactions. Science 316, 1497–1502 (2007).

[2] Vogel, M. J., Peric-Hupkes, D. & van Steensel, B. Detection of in vivo protein-DNA interactions using DamID in mammalian cells. Nat Protoc 2, 1467–1478 (2007).

[3] Giresi, P. G., Kim, J., McDaniell, R. M., Iyer, V. R. & Lieb, J. D. FAIRE (Formaldehyde-Assisted Isolation of Regulatory Elements) isolates active regulatory elements from human chromatin. Genome Res. 17, 877–885 (2007).

[4] Boyle, A. P. et al. High-resolution mapping and characterization of open chromatin across the genome. Cell 132, 311–322 (2008).

[5] Buenrostro, J. D., Giresi, P. G., Zaba, L. C., Chang, H. Y. & Greenleaf, W. J. Transposition of native chromatin for fast and sensitive epigenomic profiling of open chromatin, DNA-binding proteins and nucleosome position. Nat. Methods 10, 1213–1218 (2013).

[6] Lieberman-Aiden, E. et al. Comprehensive mapping of long-range interactions reveals folding principles of the human genome. Science 326, 289–293 (2009).

[7] Fullwood, M. J. et al. An oestrogen-receptor-alpha-bound human chromatin interactome. Nature 462, 58–64 (2009).

[8] Shiraki, T. et al. Cap analysis gene expression for high-throughput analysis of transcriptional starting point and identification of promoter usage. Proc. Natl. Acad. Sci. USA 100, 15776–15781 (2003).

[9] Davis, C. A. et al. The Encyclopedia of DNA elements (ENCODE): data portal update. Nucleic Acids Res. 46, D794–D801 (2018).

[10] Farnham, P. J. Insights from genomic profiling of transcription factors. Nat. Rev. Genet. 10, 605–616 (2009).

[11] Agrawal, A., Sambare, S. V., Narlikar, L. & Siddharthan, R. THiCweed: fast, sensitive detection of sequence features by clustering big datasets. Nucleic Acids Res. 46, e29 (2018).

[12] Eggeling, R. Disentangling transcription factor binding site complexity. Nucleic Acids Res. (2018).

[13] Staden, R. Computer methods to locate signals in nucleic acid sequences. Nucleic Acids Res. 12, 505–519 (1984).

[14] Khan, A. et al. JASPAR 2018: update of the open-access database of transcription factor binding profiles and its web framework. Nucleic Acids Res. 46, D260–D266 (2018).

[15] Bailey, T. & Elkan, C. Fitting a mixture model by expectation maximization to discover motifs in biopolymers. In Intelligent Systems for Molecular Biology, pages 28–36. AAAI Press (1994).

[16] Heinz, S. et al. Simple combinations of lineage-determining transcription factors prime cis-regulatory elements required for macrophage and B cell identities. Mol. Cell 38, 576–589 (2010).

[17] Gupta, S., Stamatoyannopoulos, J. A., Bailey, T. L. & Noble, W. S. Quantifying similarity between motifs. Genome Biol. 8, R24 (2007).

[18] Phillips, J. E. & Corces, V. G. CTCF: master weaver of the genome. Cell 137, 1194–1211 (2009).

[19] Matthews, N. E. & White, R. Chromatin Architecture in the Fly: Living without CTCF/Cohesin Loop Extrusion?: Alternating Chromatin States Provide a Basis for Domain Architecture in Drosophila. Bioessays 41, e1900048 (2019).

[20] Ni, X. et al. Adaptive evolution and the birth of CTCF binding sites in the Drosophila genome. PLoS Biol. 10, e1001420 (2012).

[21] Negre, N. et al. A cis-regulatory map of the Drosophila genome. Nature 471, 527–531 (2011).

[22] Smith, S. T. et al. Genome wide ChIP-chip analyses reveal important roles for CTCF in Drosophila genome organization. Dev. Biol. 328, 518–528 (2009).

[23] Ohler, U., Liao, G. C., Niemann, H. & Rubin, G. M. Computational analysis of core promoters in the Drosophila genome. Genome Biol. 3, RESEARCH0087.1–0087.12 (2002).

[24] Ohler, U. Identification of core promoter modules in Drosophila and their application in accurate transcription start site prediction. Nucleic Acids Res. 34, 5943–5950 (2006).

[25] Narlikar, L. Multiple novel promoter-architectures revealed by decoding the hidden heterogeneity within the genome. Nucleic Acids Res. 42, 12388–12403 (2014).

[26] Maksimenko, O. et al. Two new insulator proteins, Pita and ZIPIC, target CP190 to chromatin. Genome Res. 25, 89–99 (2015).

[27] Cuartero, S., Fresán, U., Reina, O., Planet, E. & Espinàs, M. L. Ibf1 and Ibf2 are novel CP190-interacting proteins required for insulator function. EMBO J. 33, 637–647 (2014).

[28] Narlikar, L. MuMoD: a Bayesian approach to detect multiple modes of protein-DNA binding from genome-wide ChIP data. Nucleic Acids Res. 41, 21–32 (2013).

[29] Eggeling, R. et al. On the value of intra-motif dependencies of human insulator protein CTCF. PLoS ONE 9, e85629 (2014).

[30] Heidari, N. et al. Genome-wide map of regulatory interactions in the human genome. Genome Res. 24, 1905–1917 (2014).

[31] Ye, B. et al. ZNF143 in Chromatin Looping and Gene Regulation. Front Genet 11, 338 (2020).

[32] John, S. et al. Chromatin accessibility pre-determines glucocorticoid receptor binding patterns. Nat. Genet. 43, 264–268 (2011).

[33] Biddie, S. C. et al. Transcription factor AP1 potentiates chromatin accessibility and glucocorticoid receptor binding. Mol. Cell 43, 145–155 (2011).

[34] Grøntved, L. et al. C/EBP maintains chromatin accessibility in liver and facilitates glucocorticoid receptor recruitment to steroid response elements. EMBO J. 32, 1568–1583 (2013).

[35] McDowell, I. C. et al. Glucocorticoid receptor recruits to enhancers and drives activation by motif-directed binding. Genome Res. 28, 1272–1284 (2018).

[36] Andersson, R. et al. An atlas of active enhancers across human cell types and tissues. Nature 507, 455–461 (2014).

[37] Danko, C. G. et al. Identification of active transcriptional regulatory elements from GRO-seq data. Nat. Methods 12, 433–438 (2015).

[38] Azofeifa, J. G. et al. Enhancer RNA profiling predicts transcription factor activity. Genome Res. 28, 334–344 (2018).

[39] Core, L. J. et al. Analysis of nascent RNA identifies a unified architecture of initiation regions at mammalian promoters and enhancers. Nat. Genet. 46, 1311–1320 (2014).

[40] Haeussler, M. et al. The UCSC Genome Browser database: 2019 update. Nucleic Acids Res. 47, D853–D858 (2019).

[41] Madeira, F. et al. The EMBL-EBI search and sequence analysis tools APIs in 2019. Nucleic Acids Res. 47, W636–W641 (2019).

[42] Maston, G. A., Evans, S. K. & Green, M. R. Transcriptional regulatory elements in the human genome. Annu. Rev. Genomics Hum. Genet. 7, 29–59 (2006).

[43] Heintzman, N. D. et al. Histone modifications at human enhancers reflect global cell-type-specific gene expression. Nature 459, 108–112 (2009).

[44] Cheng, J. et al. A role for H3K4 monomethylation in gene repression and partitioning of chromatin readers. Mol. Cell 53, 979–992 (2014).

[45] Liu, Y. et al. Functional assessment of human enhancer activities using whole-genome STARR-sequencing. Genome Biol. 18, 219 (2017).

[46] Lee, D. et al. Starrpeaker: Uniform processing and accurate identification of starr-seq active regions. bioRxiv (2020).

[47] Moore, J. E. et al. Expanded encyclopaedias of DNA elements in the human and mouse genomes. Nature 583, 699–710 (2020).

[48] Kim, T. H. et al. Analysis of the vertebrate insulator protein CTCF-binding sites in the human genome. Cell 128, 1231–1245 (2007).

[49] Bailey, S. D. et al. ZNF143 provides sequence specificity to secure chromatin interactions at gene promoters. Nat Commun 2, 6186 (2015).

[50] Romano, O. & Miccio, A. GATA factor transcriptional activity: Insights from genome-wide binding profiles. IUBMB Life 72, 10–26 (2020).

[51] Cusanovich, D. A. et al. The cis-regulatory dynamics of embryonic development at single-cell resolution. Nature 555, 538–542 (2018).

[52] Yanez-Cuna, J. O. et al. Dissection of thousands of cell type-specific enhancers identifies dinucleotide repeat motifs as general enhancer features. Genome Res. 24, 1147–1156 (2014).

[53] Su, Z., Wilson, B., Kumar, P. & Dutta, A. Noncanonical Roles of tRNAs: tRNA Fragments and Beyond. Annu. Rev. Genet. (2020).

[54] Sreekumar, L. et al. Orc4 spatiotemporally stabilizes centromeric chromatin. bioRxiv (2019).

[55] Arbel, H. et al. Exploiting regulatory heterogeneity to systematically identify enhancers with high accuracy. Proc. Natl. Acad. Sci. U.S.A. 116, 900–908 (2019).

[56] Hamm, D. C. & Harrison, M. M. Regulatory principles governing the maternal-to-zygotic transition: insights from Drosophila melanogaster. Open Biol 8, 180183 (2018).

[57] Reiter, F., Wienerroither, S. & Stark, A. Combinatorial function of transcription factors and cofactors. Curr. Opin. Genet. Dev. 43, 73–81 (2017).

[58] Zhou, Q. & Wong, W. H. CisModule: de novo discovery of cis-regulatory modules by hierarchical mixture modeling. Proc. Natl. Acad. Sci. U.S.A. 101, 12114–12119 (2004).

[59] Xie, D. et al. Dynamic trans-acting factor colocalization in human cells. Cell 155, 713–724 (2013).

[60] Giannopoulou, E. G. & Elemento, O. Inferring chromatin-bound protein complexes from genome-wide binding assays. Genome Res. 23, 1295–1306 (2013).

[61] Guo, Y. & Gifford, D. K. Modular combinatorial binding among human trans-acting factors reveals direct and indirect factor binding. BMC Genomics 18, 45 (2017).

[62] Benton, M. L., Talipineni, S. C., Kostka, D. & Capra, J. A. Genome-wide enhancer annotations differ significantly in genomic distribution, evolution, and function. BMC Genomics 20, 511 (2019).

[63] Bailey, T. L. & Machanick, P. Inferring direct DNA binding from ChIP-seq. Nucleic Acids Res. 40, e128 (2012).

[64] Liu, J. The collapsed gibbs sampler with applications to a gene regulation problem. J. Am. Stat. Assoc. 89, 958–966 (1994).

[65] Mitra, S., Biswas, A. & Narlikar, L. DIVERSITY in binding, regulation, and evolution revealed from high-throughput ChIP. PLoS Comput. Biol. 14, e1006090 (2018).

